# Germline *de novo* mutation rates on exons versus introns in humans

**DOI:** 10.1101/2019.12.23.886879

**Authors:** Miguel Rodriguez-Galindo, Sònia Casillas, Donate Weghorn, Antonio Barbadilla

## Abstract

A main assumption of molecular population genetics is that genomic mutation rate does not depend on sequence function. Challenging this assumption, a recent study has found a reduction in the mutation rate in exons compared to introns in somatic cells. This reduction was ascribed to an enhanced exonic mismatch repair system activity. If this reduction happens also in the germline, it can compromise studies of population genomics, including the detection of the footprint of selection when using introns as proxies of neutrality. Here we compiled and analyzed published germline *de novo* mutation (DNM) data to test if the exonic mutation rate is also reduced in germ cells. We detected ascertainment bias in studies using DNM data from diseased probands and investigated the impact of extended nucleotide context on *de novo* mutation rate. After controlling for these factors, we found no reduction in the mutation rate in exons compared to introns in the germline genome, in contrast to what has been previously described in somatic cells. Therefore, there is no evidence of an enhanced mismatch repair system activity in exons with respect to adjacent introns in germline cells.

---

One of the most general and widely accepted predictions of the neutral theory of molecular evolution is that “the more sequence conservation, the more functional (selective) constraint on the sequence” [1]. This principle explains why different functional regions in the genome have different levels of polymorphism and divergence, such as the lower variation at nonsynonymous vs synonymous sites in protein-coding genes or in exonic vs intronic sequences [2]. This relationship between constraint and variation constitutes one of the most powerful approaches in the current search for functional regions in the genome and the detection of natural selection at the molecular level. An integral part of estimating constraint, or purifying selection, on functional genomic regions is the comparison of the observed number of mutations to the expectation under neutral evolution. In genes, this neutral expectation is usually estimated from putatively non-functional regions or sites, including intronic sequence [3, 4]. A main requirement for the validation of this assumption is that mutation rate on exons and introns does not correlate with that sequence function.

Mutation rates can vary strongly across the human genome, with positional differences up to 3-fold in the germline [5] and up to 5-fold in tumor cells [6]. It is influenced by several factors, including replication time, chromatin state, and expression level [7, 8, 6]. A priori, none of these factors are expected to correlate directly with genic sequence function (exonic vs intronic). Another important determinant of mutation rate is DNA sequence composition. Recent studies have addressed mutational processes and their associated sequence-dependent “signatures”, both in the soma and the germline [9, 10, 11, 12]. Germline and many cancer tumor signatures exhibit a higher relative rate of C>T transitions for single nucleotide variants (SNV) [13, 9, 6]. Consequently, due to their higher G/C content, exonic regions show a context-driven relative increase in mutation rate compared to intronic regions [14], which can be corrected for with the proper mutational model.

A differential mutation rate between intronic and exonic DNA beyond the context dependence would require a mutational process that recognizes the difference between the two functional sequence categories. Recently, Frigola et al. [15] found in tumoral DNA, primarily from skin melanomas and POLE-aberrant colorectal cancers, that mutation rates are surprisingly lower in exons than in introns after accounting for the trinucleotide-context-dependent mutational signature. This reduced mutation rate in exons is similar both in synonymous and nonsynonymous sites, which rules out purifying selection as an explanation. The study suggests that the lower mutation rate in exons results from an enhanced mismatch repair (MMR) activity in exons compared to introns. In turn, the increased repair activity is attributed to differential amounts of H3K36me3 epigenetic marks of exons and introns. If this observation of different mutation rates between introns and exons could be extrapolated to the germline, as the authors suggest, then population and functional genomics studies would be compromised, and they should include these differential mutation rates as an integral part of their explanatory models.

Here, we investigated the relative mutation rates of exons and introns in the human germline using *de novo* mutation (DNM) data. We aggregated 679,547 SNV DNMs from 7 family-based WGS datasets, consolidating a high-density, high-quality DNM map across the human genome (**Methods, Supplementary Figure S1**). We first analyzed whether exonic and intronic mutation densities differ among human DNMs after accounting for sequence composition. We computed the observed total mutation burden at exonic and intronic sites by summing over 95,633 internal exon-centered sequences of size 2,001 base pairs (bp), carrying a subset of 50,780 genic mutations. We compared this observation to the expected mutation burden on exons and introns. Since per-nucleotide mutation probability is influenced by the neighboring sequence context, we derived the expected mutation burden at each position of each 2,001-bp internal exon-centered window from a context-dependent model.

We first used a trinucleotide-context-dependent germline whole-genome mutation signature model, in line with the analysis presented in Frigola et al. [15] for somatic mutations. That study found that the mutation burden of POLE-mutant tumors in positions dominated by exonic DNA is lower than expected (**Figure 1a**). In contrast, the observed germline exonic mutation burden was significantly increased by 7.2% (s.d. 1.4%; P=0.001, permutation-based test) compared to the expectation across introns and exons (**Figure 1b**). This result is robust to biases in mutation calling due to region mappability differences (**Supplementary Table S1, Supplementary Figure S2**) and effects of transcription-coupled repair on the mutational signature (**Supplementary Figure S3**). This suggests that the hypothesized mechanism of enhanced repair on exons compared to introns in POLE-deficient tumors is not determining mutation rate in the germline.

**Figure 1.**
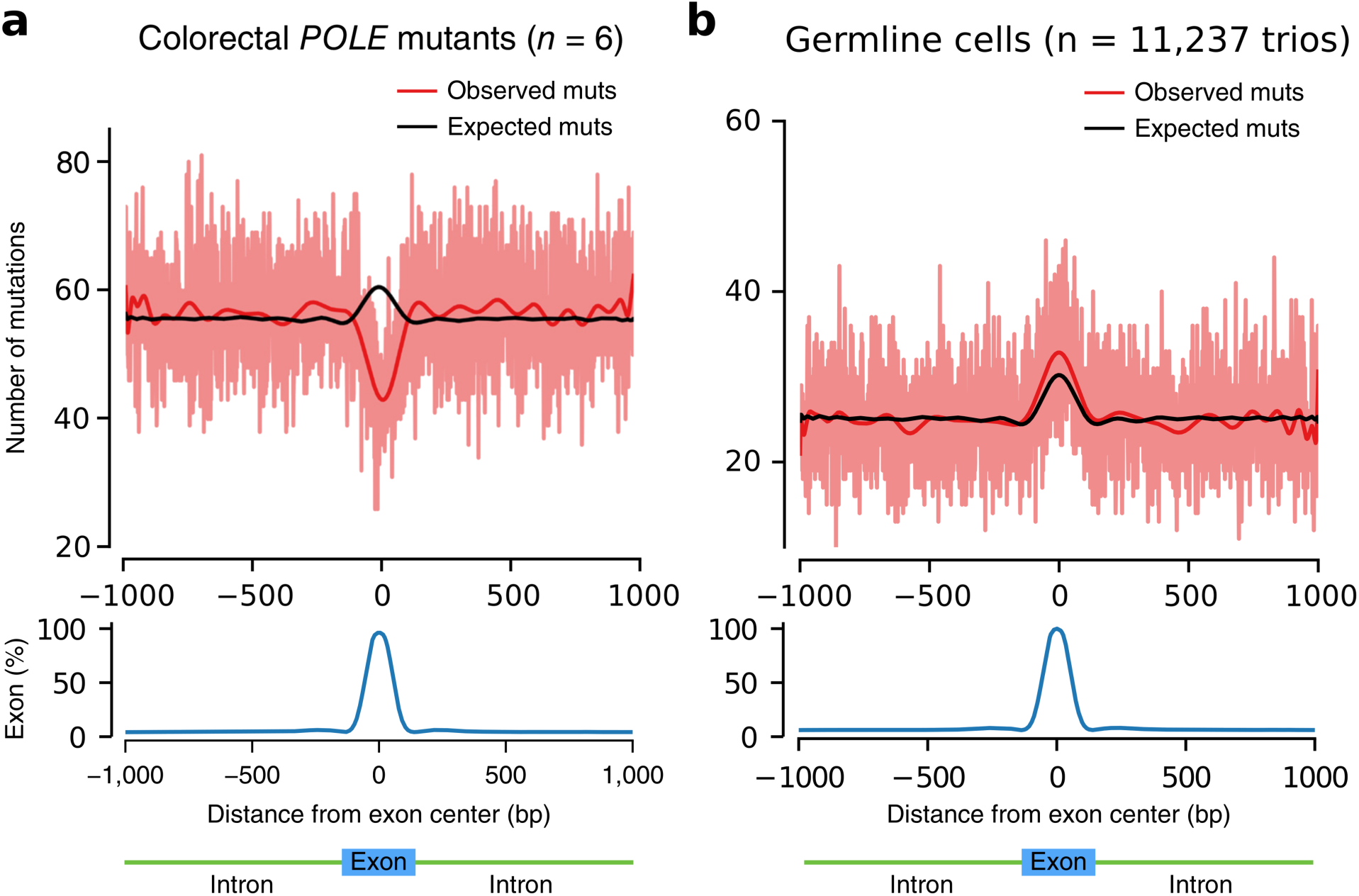
Internal exon-centered analyses on somatic and germline *de novo* mutations. Exon-centered 2,001-nt-wide observed and expected mutational profiles (top) and exon density (bottom) in (a) somatic and (b) germline cells. The light red line represents the observed number of mutations at each position, whereas the dark red and black lines represent smoothed numbers of observed and expected mutations, respectively, obtained from a polynomial fit. **(a)** Profile of mutations in six POLE-mutant colorectal tumors, Figure 2a from Frigola et al. [15]. **(b)** Profile of mutations in the germline of 11,237 trios.

Even in the absence of the supposed enhanced MMR effect on exonic DNA, we would expect a very slight deficit of exonic relative to intronic DNMs, due to strong purifying selection on lethal *de novo* variants in the early stages of embryonic development [16]. Therefore, we next investigated potential factors that could explain the observed increase in exonic relative to intronic mutation burden in DNM datasets, such as technical differences in sequencing and calling (**Supplementary Table S2**) or enrichment in diseased probands. As detailed in **Table 1**, the analyzed DNM datasets are heterogeneous regarding their study conditions, including disease cohorts. Diseased probands, e.g. those with autism spectrum disorder (ASD) or preterm birth, are more likely than average to carry mutations with functional impact [17, 18]. This ascertainment bias in the data is expected to entail an enrichment with exonic nonsynonymous variants [19, 20]. To test this, we classified the 4,669 mutations in the internal exons as synonymous and nonsynonymous changes, resulting in 3,488 nonsynonymous and 1,170 synonymous DNMs (corresponding to a ratio of 2.98:1). We then repeated the internal exon-centered analysis for each of the two exonic mutation categories.

**Table 1.**
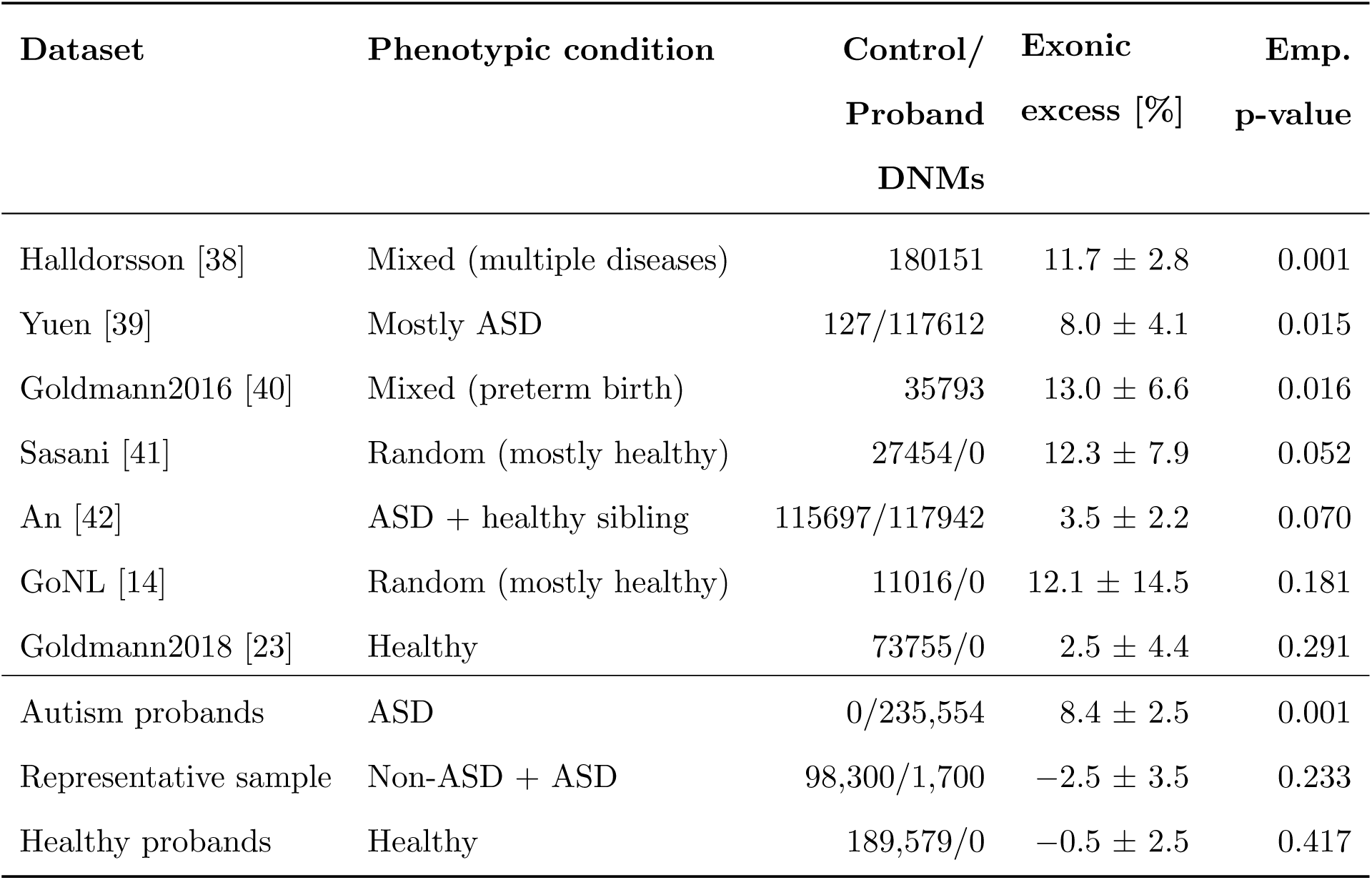
Properties of analyzed DNM datasets, including excess in exonic burden. The seven used datasets and their references are shown above the line, while below the results for the composed datasets (see **Methods**) are given. In each group, datasets are ordered from most to least significant exonic mutation excess. Errors denote one s.d.

**Figure 2a** shows that the observed synonymous profile matches the expected profile almost perfectly, with a slight non-significant deficit (−1.1%, P=0.353). However, exonic nonsynonymous mutations show a large and statistically significant excess compared to the expectation under the trinucleotide context model (10.4%, P=0.001, **Figure 2b**). Note that this stratification by functional mutation category entails a reduction of the number of both synonymous and nonsynonymous mutations relative to flanking introns across the window of stacked sequences. This is why the number of mutations, when moving outwards from the center, converges to that of **Figure 1b**. Overall, **Figure 2** suggests that disease ascertainment during data acquisition may be responsible for the overall excess of 7.2% of exonic variants. Therefore, we next repeated the analysis for all seven DNM datasets individually, as well as for assembled samples with only healthy, only ASD, or a representative mixture of probands (**Methods**). We found a significant exonic mutation excess only in mixed cohorts or those assembled purely from diseased samples, while all cohorts with mostly or exclusively healthy probands show no signal (**Table 1**). Moreover, when we stratify exonic mutations by functional impact, we find a statistically significant excess only in nonsynonymous mutations among ASD individuals (**Supplementary Table S3**).

**Figure 2.**
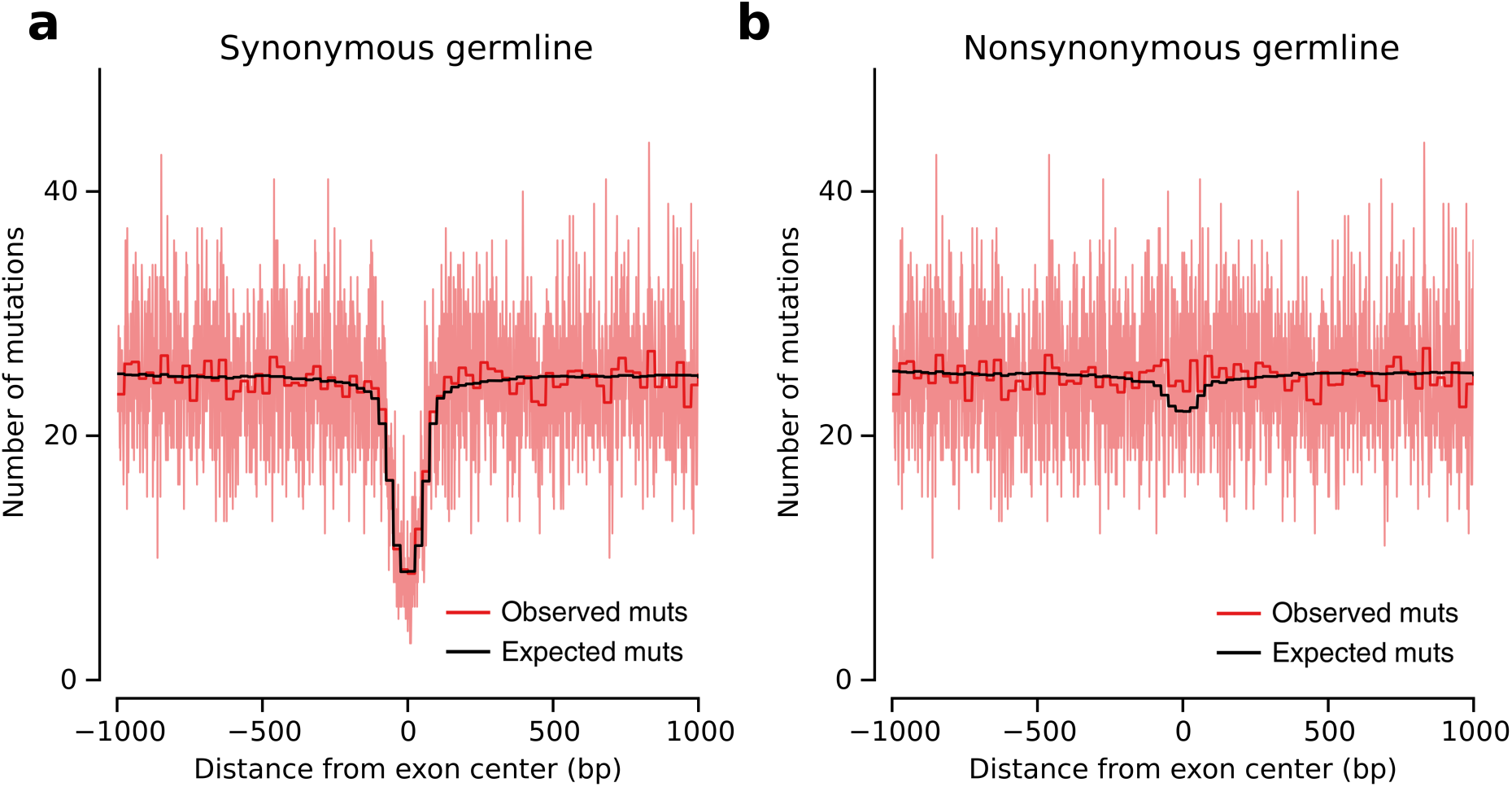
Internal exon-centered analysis for synonymous and nonsynonymous DNMs. The light red line represents the observed number of mutations at each nucleotide position, while the dark red and black lines represent averages in bins of size 25 positions for observed and expected mutations, respectively. **(a)** Synonymous DNM profile with observed exonic mutation difference of −1.1% (P=0.353). **(b)** Nonsynonymous DNM profile, showing an exonic excess of 10.4% (P=0.001). Due to the removal of nonsynonymous and synonymous mutations in panels (a) and (b), respectively, the total number of exonic mutations relative to flanking introns is reduced. The number of mutations, when moving away from the center, converges to that of Figure 1b.

The stratification into synonymous and nonsynonymous changes entailed a polarization of the exonic excess (**Figure 2**), intensifying the signal for nonsynonymous variants (10.4%) with a concomitant decrease for synonymous variants (−1.1%). This type of polarization could be due to the incompleteness of our mutational model. Our mutational model for the expected number of mutations was constructed using trinucleotide-context-dependent mutation probabilities. However, it has been shown that SNPs segregating in the human population are affected by the extended flanking sequence, with a heptameric context explaining a majority of the observed mutation rate variability [21, 12]. **Figure 3** shows that this is confirmed by DNMs based on the relative frequencies of all four nucleotides around mutations in our DNM dataset, although the effect is small compared to the one observed in POLE-mutated tumor genomes [22].

**Figure 3.**
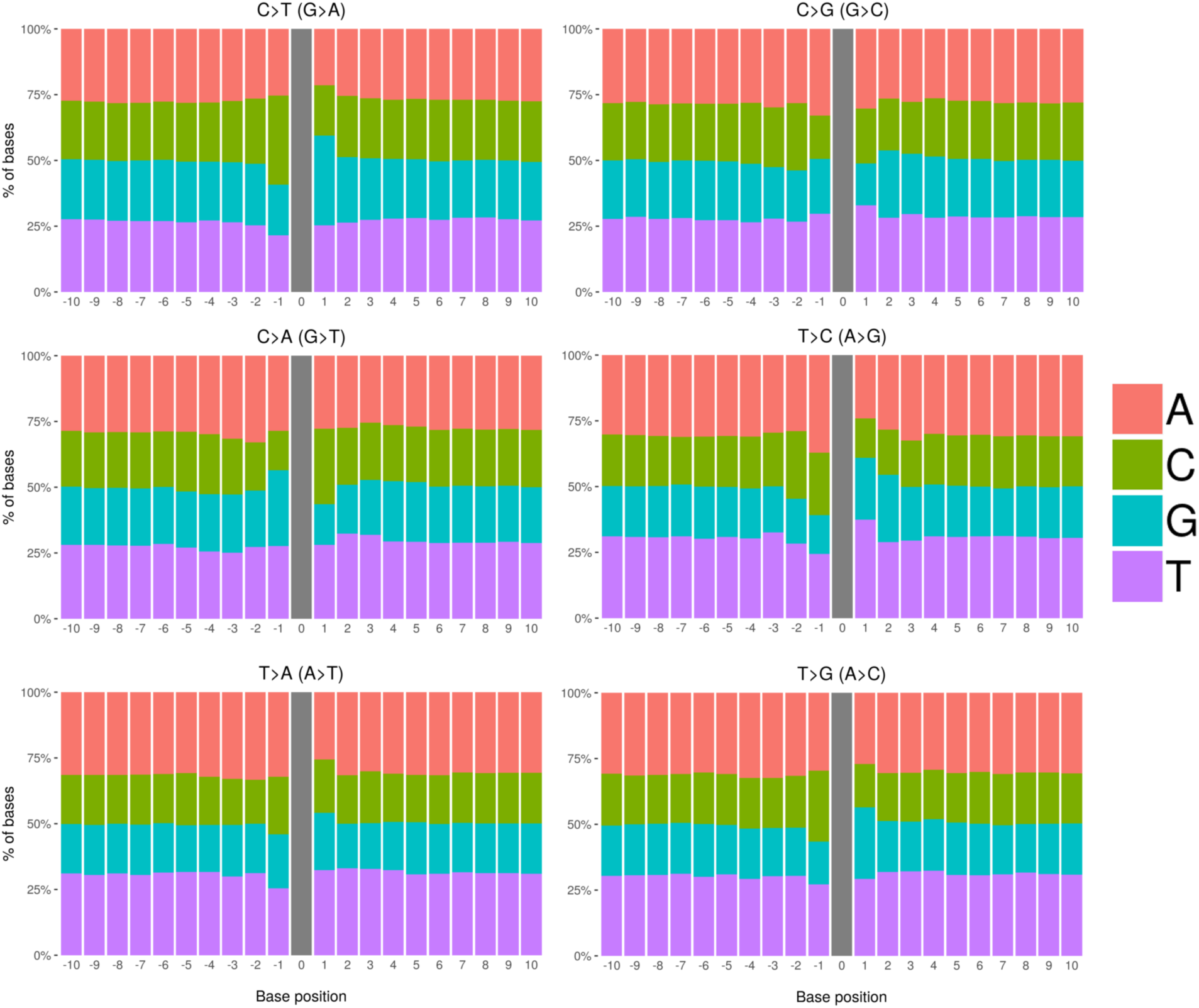
Sequence context around *de novo* mutations. Frequencies of nucleotides neighboring our set of 679,547 DNMs in a window of size 21 bp across each 1-mer mutation class. The amount of extended sequence context dependency varies across mutation classes.

We assessed the impact of context dependency by expanding our mutational signature model to incorporate the pentameric and heptameric mutational sequence context (based on exact computation and a likelihood decomposition approach, respectively; **Methods, Supplementary Figure S4**). We applied these extended-context models to the largest DNM dataset that had no diseased probands (Goldmann et al., 2018 [23]), as well as to this dataset and the pooled dataset stratified by synonymous and nonsynonymous variants. We found that while the overall likelihood increases for increasing context size, penalization due to the additional parameters of the larger context models entails that the trinucleotide-context-dependent model is found to be the best model for the current datasets based on the Akaike information criterion (AIC) (**Table 2, Supplementary Tables S4** and **S5**).

**Table 2.** Extended sequence context dependency for Goldmann et al. (2018). Errors denote one s.d.

| Model | % increase | Emp. p-value | Log-likelihood | # param | AIC |
| --- | --- | --- | --- | --- | --- |
| 1-mer | $16.2 \pm 5.4$ | 0.001 | -68111.4 | 12 | 136246.8 |
| CpG | $1.7 \pm 4.3$ | 0.348 | -66594.1 | 18 | 133224.2 |
| 3-mer | $2.7 \pm 4.3$ | 0.281 | -66315.0 | 192 | 133014.1 |
| 5-mer | $2.9 \pm 4.6$ | 0.258 | -66054.5 | 3072 | 138253.0 |
| 7-mer | $2.8 \pm 4.3$ | 0.266 | -65981.7 | 49152 | 230267.5 |

Whether originating from replication errors, spontaneous deaminations or oxidative damage, the dominant mutational processes in the germline are expected to produce mismatches [24, 25, 26, 27, 28]. Thus, the MMR mechanism is expected to play a key role in the repair of germline cells. We therefore investigated whether this mechanism is exonically enhanced in the germline, as was proposed for POLE-aberrant tumors via the H3K36me3 mark [15]. Using our dataset of mutations from healthy probands, we evaluated the density of H3K36me3 marks and nucleosomes. These two features were reported to show differential coverage between (mainly internal) exons and introns (**Supplementary Figure S5**), and had been previously described to contribute to the recognition of splice marks at internal exon-intron boundaries [29, 30]. We observed no significant correlation (*r* = −0.03, *P* = 0.84) between the exonic mutation enrichment and the H3K36me3 mark (**Figure 4b**). This is in contrast to the recruitment mechanism described in somatic cells [31], which is invoked as mechanistic hypothesis behind the results of the study by Frigola et al. [15] (**Figure 4a**). Conversely, we find a negative, nearly significant correlation (*r* = −0.28, *P* = 5.38 × 10^−2^) with nucleosome coverage (**Supplementary Figure S6**). This could be explained by replication errors being preferentially repaired at positioned nucleosomes, in line with the post-replicative nucleosome-reconstitution-coupled MMR activity [32].

**Figure 4.**
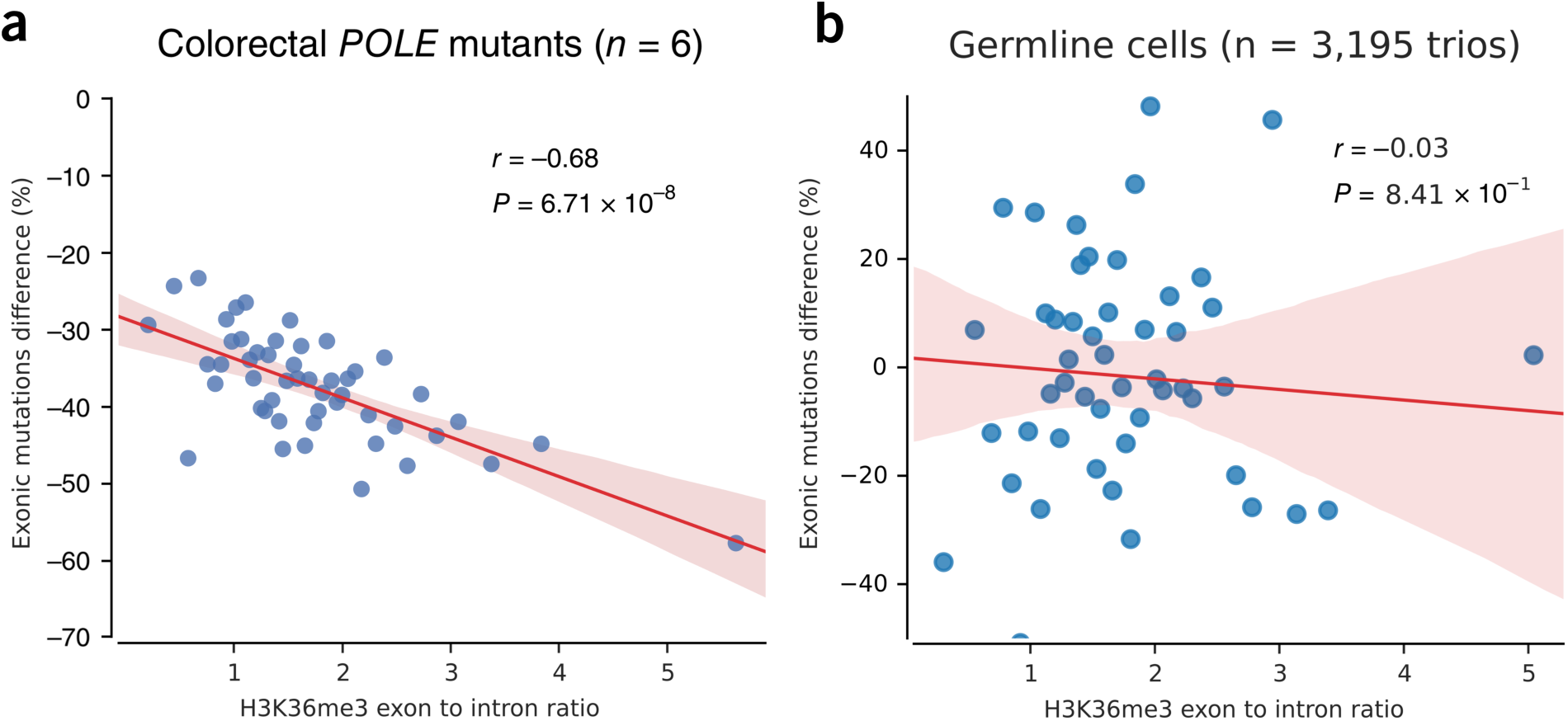
Deviation in the exonic mutation burden as a function of the H3K36me3 exon-to-intron ratio. Blue dots denote 50 groups of genes binned by their exon-to-intron ratio of H3K36me3 coverage (x-axis). The relative difference between the total observed and expected number of exonic mutations (computed using a 3-mer model) per group is shown on the y-axis. The trendline and its confidence interval were added using the seaborn package of Python, while the correlation coefficient and its significance were computed using an iteratively re-weighted least-squares approach. **(a)** POLE-mutant colorectal tumors. The H3K36me3 histone mark is derived from colonic mucosa (E075). Modified from Frigola et al. [15]. **(b)** DNMs from healthy probands (a total of 3,195 trios). The H3K36me3 histone mark is derived from H1 stem cells (E003).

In summary, using recent aggregated human *de novo* mutation data we showed that the rate of generation of new genetic variants, the mutation rate, does not significantly vary between functionally distinct genic regions in the germline genome. This finding supports the validity of the selective constraint assumption when searching for functional regions in the genome and the footprint of natural selection at the molecular level. Therefore, our analysis shows that the results found in the soma by Frigola et al. [15] cannot be directly extrapolated to the germline, and the MMR-dependent process that was proposed as an explanation for the decreased exonic mutation rates in somatic cells does not seem to determine germinal cell mutation rates.

The mutational process in somatic cells is different from the one in germinal cells. In mammals, the spontaneous mutation rate in germline is lower than in somatic cells [33, 16], a difference of more than one order of magnitude in humans [26]. Furthermore, tumors greatly vary in their mutation spectrum [34, 9, 35], reflecting particular differences among cell types in repair and mutagenic mechanisms, while germline variation is mainly driven by CpG transitions [36]. Our estimates of exonic and intronic mutation rate of, respectively, 1.38 × 10^−8^ and 1.11 × 10^−8^ mutations per site per generation, obtained from the representative Goldmann dataset [23], are consistent with the previously reported whole-genome based rate of 1.2 × 10^−8^ estimated by Millholland et al. [26]. Estimates obtained from the pooled dataset reflect the disease ascertainment bias, with a similar intronic mutation rate (1.15 × 10^−8^), but a much larger exonic rate (1.52 × 10^−8^), in line with previous findings in diseased cohorts [37].

Through aggregating *de novo* data across cohorts to describe germline mutation patterns, we went beyond previous analyses which used extreme rare variants as a proxy for DNMs [21, 12]. We corroborate earlier findings, in particular the heptameric context dependence of germline variants. At the same time, the internal exon-centered analysis with its relatively low number of mutations is still adequately described by a trinucleotide-context-dependent model. Beyond context dependence, the ascertainment bias introduced by enrichment with diseased probands is one of the most important confounding factors of DNM analyses. We showed that its effects can lead to significant deviations from the null model and, depending on the application, should be addressed through an informed choice of samples. This study provides a clear-cut answer to the challenge posed by Frigola et al. [15]: it validates the main assumption of molecular population genetics and points out different mutational dynamics of somatic vs germinal cells.

## Acknowledgments

We thank especially Núria López-Bigas for her generosity, thoughtful insight and suggestions through the numerous discussions during the development of this work. We also thank her research group for sharing the software code in all their contributions, especially their Frigola et al. (2017) paper. The project that gave rise to these results received the support of a fellowship from “la Caixa” Foundation (ID 100010434) with fellowship code LCF/BQ/DR19/11740019 [M.R.]; by the Ministerio de Economía y Competitividad (Spain) [CGL2017-89160P to M.R. and A.B.]; and AGAUR (Generalitat de Catalunya) [2017SGR-1379 to A.B.].

## Data availability

Mutation datasets were downloaded from the supplementary tables of the respective papers (Goldmann et al., 2016; Halldorsson et al., 2019; Yuen et al., 2017; An et al., 2018; Sasani et al., 2019) or by direct request to the authors (Goldmann et al., 2018). Data from the GoNL was downloaded from http://www.nlgenome.nl. We downloaded narrow peak coordinates and genome-wide read-coverage of H3K36me3 from human embryonic stem cell H1-hESC (E003) from the Epigenome Roadmap consortium. The genome-wide nucleosome positioning density graph of ENCODE cell line GM12878 (lymphoblastoid cell line) was obtained via the UCSC genome browser http://hgdownload.cse.ucsc.edu. Coordinates of exonic and intronic boundaries after mappability filtering can be obtained from the authors upon request.

## Code availability

Whenever possible, code published in Frigola et al. (2017) was used in pure or slightly adapted form to ensure comparability. Custom scripts for data analysis were used according to the descriptions in the Methods section and are available upon request from the authors.

## Methods

### *De novo* mutation data

We aggregated DNMs from 7 family-based WGS datasets coming from multiple centers and projects: the Genomes of the Netherlands (GoNL) project [14], the Inova Translational Medicine Institute Preterm Birth Study [40], Inova Translational Medicine Institute’s Longitudinal Childhood Genome Study [23], deCODE genetics [38], Simons Simplex Collection (SCC) and Korean ASD cohort [42], the Autism Genetic Research Exchange (AGRE) repository [39] and Centre d’Etude du Polymorphisme Humain (CEPH) [41]. Mutation datasets were downloaded from the supplementary tables of the respective papers [40, 38, 39, 42, 41] or by direct request to the authors [23]. Data from the GoNL was downloaded from http://www.nlgenome.nl. Most of datasets were originally mapped to hg19, with exception of the data from An et al. 2018 [42] and Halldorson et al. 2019 [38], which were mapped to hg38. Subsequently, coordinates in these datasets were lifted over to hg19, the most common reference genome in our data. To avoid possible biases arising from mutation calling on sexual chromosomes, only autosomal SNVs were used, leaving a total of 679,547 germline SNV DNMs coming from 11,237 trios.

### Effect prediction of *de novo* mutations

The predicted consequence class of all DNMs was obtained using the Ensembl Variant Effect Predictor (VEP) [43] for the GRCh37/hg19 assembly. Since some DNMs were reported to have more than one consequence, e.g. different transcripts or overlapping genes, only one predicted consequence for each DNM was retrieved (according to VEP criteria). Predictions are classified from major to mild according to the Ensembl Variation hierarchy.

### Whole genome analysis of the *de novo* mutation spectrum

All mutations were divided into 9 classes, considering the fact that CpG sites are highly mutagenic. The number of mutations were corrected by the relative abundance of the context in the whole genome, e.g. the total number of C>T (G>A) transitions occurring at CpG sites divided by the relative abundance of CpG sites in the genome. We performed the mutational analysis across all used studies (**Supplementary Figure S1**). For all subsequent analyses, extended nucleotide-context-dependent mutational models were used.

### Genomic coordinates of internal exons and flanking introns

Coordinates for a total of 20,345 protein-coding genes were obtained from GENCODE v19 [44]. Genes without introns and overlapping genes were discarded, leaving a filtered set of 13,474 genes. Genes located on chromosomes X, Y and on the mitochondrial genome were removed from the analysis, leaving a total of 12,754 autosomal genes. Finally, all transcripts per gene were merged into meta-exon and meta-intron coordinates, both 5’ and 3’ flanking exons were removed as well as UTRs (**Supplementary Tables S6, S7**). Only internal exons (unfiltered by mappability issues, see below) were used for the main internal exon-centered mutational analyses.

Moreover, positions where mutation calling would be technically challenging because of mappability issues were removed, leaving a total of 10,237 genes for the gene by gene analyses. We also filtered out the internal exon-centered 2001-nt windows that overlapped at least one nucleotide with regions with mappability issues (**Supplementary Table S1**) for the supplementary internal exon-centered mutational analysis (**Supplementary Figure S2**). Coordinates of unreliable regions were obtained from the UCSC Genome Browser, available at http://genome.ucsc.edu/cgi-bin/hgFileUi?db=hg19&g=wgEncodeMapability.

Meta-exon and meta-intron coordinates of genes with at least 5 meta-exons (not only internal) were extracted from GENCODE v19 for the analyses with chromatin features across genic regions (**Supplementary Figure S5**).

### Sequence context model

For the 1-mer, 1-mer with CpG, 3-mer, 5-mer and 7-mer models we directly computed the probability that a given reference single nucleotide *H*_*r*_ changes to an alternate one, *H*_*a*_ where *a* ∈ {1, 2, 3}, given its flanking sequence *X* = (*X*_5′_, *X*_3′_), where *X*_5′_ and *X*_3′_ are the 5’ and 3’ flanking sequences, respectively. Here, *H*_*r*_ ∈ *M* = {A, C, G, T}, where the latter denote nucleotides adenine, cytosine, guanine and thymine, and *H*_*a*_ ∈ *M′* = {*m* ∈ *M* | *m* ≠ *H*_*r*_}. For example, for the 5-mer GCACG > GCTCG mutation *H*_*r*_ = A, *H*_*a*_ = T, *X*_5′_ = (G, C), *X*_3′_ = (C, G) and *X* = (G, C, C, G). Therefore, the probability of each of the possible k-mer changes, normalized by the abundance of each reference k-mer in the genome, was computed as follows:

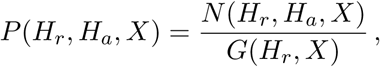

where *N* (*H*_*r*_, *H*_*a*_, *X*) is the genome-wide number of observed mutations with given flanking sequence *X* and the reference and alternate alleles *H*_*r*_ and *H*_*a*_, respectively. *G*(*H*_*r*_, *X*) is the abundance of the reference k-mer with reference nucleotide *H*_*r*_ and flanking sequence *X* in the genome. We computed the relative abundance of each reference k-mer in the autosomal genome using the pyFasta package. We also computed signatures restricted to mutations falling in genic regions (exons at the canonical CDS and the respective introns) for the supplementary analysis in **Supplementary Figure S3**.

For some k-mer models, given limitations imposed by the amount of *de novo* mutations, we used a decomposition approach to compute the probability. For a k-mer model of sequence length *k*, let *H*_*r*_ be a reference core h-mer and *H*_*a*_ an alternate core h-mer, where 1 ≤ *h* < *k*. Let the tuple *X* = (*x*_1_, …, *x*_*g*_) with *g* = (*k* − *h*) elements again represent the flanking sequence of the core h-mer, where *x*_*i*_ ∈ *M* ∀*i* ∈ {1, …, *g*}. In other words, *X* ∈ *M*^*g*^ where *M*^*g*^ is the g-fold Cartesian product. For example, with *k* = 7, *h* = 3 and the mutation ACTGACT > ACTCACT, then *H*_*r*_ = TGA, *H*_*a*_ = TCA and *X* = (*x*_1_ = A, *x*_2_ = C, *x*_3_ = C, *x*_4_ = T). We then approximate the probability *P* (*H*_*r*_, *H*_*a*_, *X*) by:

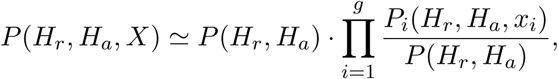

with

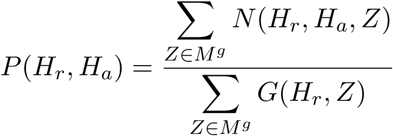

and

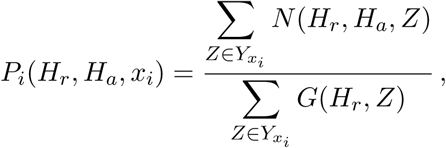

where

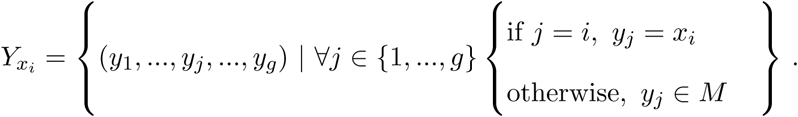

Here, *N* (*H*_*r*_, *H*_*a*_, *Z*) denotes the number of mutations in our sample with reference core *H*_*r*_, alternate core *H*_*a*_ and flanking sequence *Z*, and *G*(*H*_*r*_, *Z*) is the abundance in the genome of the k-mer with reference core *H*_*r*_ and flanking sequence *Z*.

We computed probabilities using custom Python code and applied the composite likelihood model to the 7-mer analysis of the data pooled across all cohorts using *k* = 7 and *h* = 5. For Goldmann et al. (2018), we used it for 5-mers with *k* = 5 and *h* = 3 and for 7-mers with *k* = 7 and *h* = 3. **Supplementary Figure S4** shows the relationship between the exact computations of mutational probabilities and the composite likelihood model.

### Internal exon-centered mutational analysis

A total of 95,633 stacked 2,001-nt sequences centered on the middle position of internal meta-exons were used to compare the observed and expected mutational profiles across exons and introns. We computed the frequency of mutation at a site *l* with reference core sequence 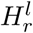 and flanking sequence *X*^*l*^ as

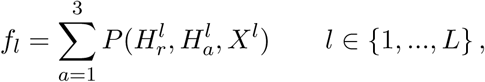

where *L* = 2001 denotes the total number of considered sites. Then, each frequency was normalized by the total frequency on the sequence:

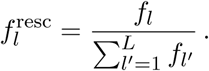

Finally, the total number of observed mutations *n*_*s*_ on each 2,001-nt sequence *s* was redistributed across both middle exonic and flanking intronic sites according to the normalized frequencies:

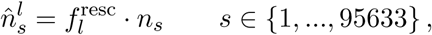

thus yielding the expected number of mutations at site *l* of a given sequence *s*. By adding up the values of all the stacked sequences we obtain the cumulative number of expected mutations at site *l*,

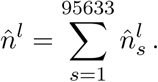

For the internal exon-centered analysis on synonymous or nonsynonymous mutations, we separated all possible exonic mutations in middle exon sequences into two groups: those with synonymous consequence and those with a consequence ranking higher than synonymous in the Ensembl Variation hierarchy. Then we computed the expected numbers by only adding frequencies for either synonymous or nonsynonymous mutations.

### Computation of effect size and statistical significance

We performed 1,000 random permutations of the observed mutations in each stacked sequence based on the probability of each site to acquire a mutation. The effect size, defined as the relative increase or decrease in observed exonic mutations with respect to the expected number, was computed based on the simulation mean expected value. The error of this estimate is given as one standard deviation derived from the 1,000 permutations. Moreover, we computed an empirical p-value as the fraction of the simulations with fewer (or more) exonic mutations than the observed number of exonic mutations.

### Composed datasets

Mutations were only resampled from datasets with known conditions of the probands, either from healthy probands or those with ASD. Given an ASD prevalence in humans of 1.7%, we created a random sample of 100,000 whole-genome DNMs, taking 98,300 mutations classified as strictly from healthy probands and 1,700 classified as ASD and repeated the internal exon-centered analysis. To generate the “healthy” and purely ASD cohorts, we used solely mutations from the respective probands.

### Nucleosome and H3K36me3 histone mark data

We downloaded narrow peak coordinates and genome-wide read-coverage of H3K36me3 from human embryonic stem cell H1-hESC (E003), as proxy for germline cells, from the Epigenome Roadmap consortium [45]. The genome-wide nucleosome positioning density graph of ENCODE [46] cell line GM12878 (lymphoblastoid cell line) was obtained via the UCSC genome browser http://hgdownload.cse.ucsc.edu. Nucleosome peak regions were identified across the genome by using the bwtool program (with parameters local-extrema -maxima -min-sep=150). 146bp flanking the peak coordinate (73bp per side) was considered the region covered by a nucleosome.

### Differential coverage of chromatin features across exons and introns

Exons and introns in each gene were classified according to their position with respect to the transcription start site, where the ones that occupy different positions in different transcripts were discarded. We also discarded exons and introns at the lower quartile of length to compute the coverage for a set of exons or introns of heterogeneous lenghts in a given position: the fraction of bases covered by H3K36me3 and nucleosomes at the center of the stack corresponding to the window defined by the shortest exon or intron remaining after the filtering. Finally, the difference between the exonic and intronic coverage was computed via the two-tailed Mann-Whitney p-value of the comparison of both distributions.

We also computed the positions in the genome covered by H3K36me3 or nucleosomes across 95,633 internal exon-centered 4,001-nt windows. By stacking sequences we obtained middle exon-centered profiles of coverage across exons and introns (**Supplementary Figure S5**).

### Nucleosome and H3K36me3 binned gene analysis

For each gene, we computed the readcount-based exonic enrichment of H3K36me3 or nucleosomes as the ratio between the exonic and intronic total number of bases covered by reads of the chromatin feature. Genes with no exonic and intronic bases coverered by reads were removed from the analysis, as well as genes without any observed exonic or intronic mutation. Thus, a total of *T* = 7215 and *T* = 6529 genes remained for the H3K36me3 and nucleosome analysis, respectively.

For a given gene, we computed the exonic expected number of mutations as follows:

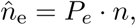

where *n* is the total number of mutations (both exonic and intronic) observed on the gene and *P*_e_ is the (binomial) probability of a mutation to fall on the exonic region of the gene, which in turn is computed as:

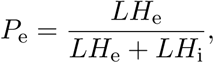

with

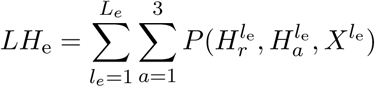

and

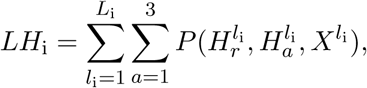

Here, *l*_e_ ∈ {1, …, *L*_e_} and *l*_i_ ∈ {1, …, *L*_i_} denotes the set of all exonic and intronic positions of a given gene, respectively. *LH*_e_ and *LH*_i_ represent the likelihood of a mutation happening in an exon or an intron, expressed as the sum of the probability 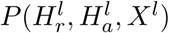 of all possible three mutations that can happen across all exonic or intronic sites of the gene, respectively. The probability was computed for each of the *T* genes as explained above only with mutations from the “healthy” composed dataset and under a 3-mer model.

Afterwards, genes were grouped into 50 bins according to their exonic enrichment of H3K36me3 or nucleosomes. Then, with the observed *n*_e_ and expected 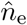 exonic mutations over all genes in the bin, we computed the relative difference between the observed and expected number of exonic mutations per bin as follows:

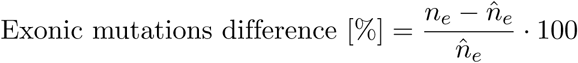

Finally, we computed the correlation between the median exonic chromatin feature enrichment and the difference in exonic mutations across the bins. The trend line and its confidence intervals were added using the bootstrapping functions of the python seaborn package, which confers equivalent weights in the regression to all points. In order to guarantee that the trend is not the result of a few outliers, the correlation coefficient and its significance were computed using an iteratively re-weighted least squares approach, letting the variance of exonic chromatin feature enrichment of the bins influence the weight of each point.

### Estimation of absolute mutation rates

We estimated absolute mutation rate in our set of 95,633 middle exons as a proxy of mean exonic mutation rate and absolute mutation rate on the rest of the 2,001-nt window as a proxy of mean intronic mutation rate as follows:

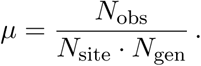

Here, *µ* is the mutation rate per site and generation, *N*_obs_ is the number of observed mutations, *N*_site_ is the number of sites (in our case 13,632,264 exonic sites and 177,729,369 flanking intronic sites) and *N*_gen_ is the number of generations. For the largest healthy dataset (Goldmann et al. 2018) [23], *N*_gen_ = 2,582 gametogeneses, while *N*_gen_ = 22,474 gametogeneses in the pooled dataset.

## Supporting Information

### Supplementary Figures

**Figure S1.**
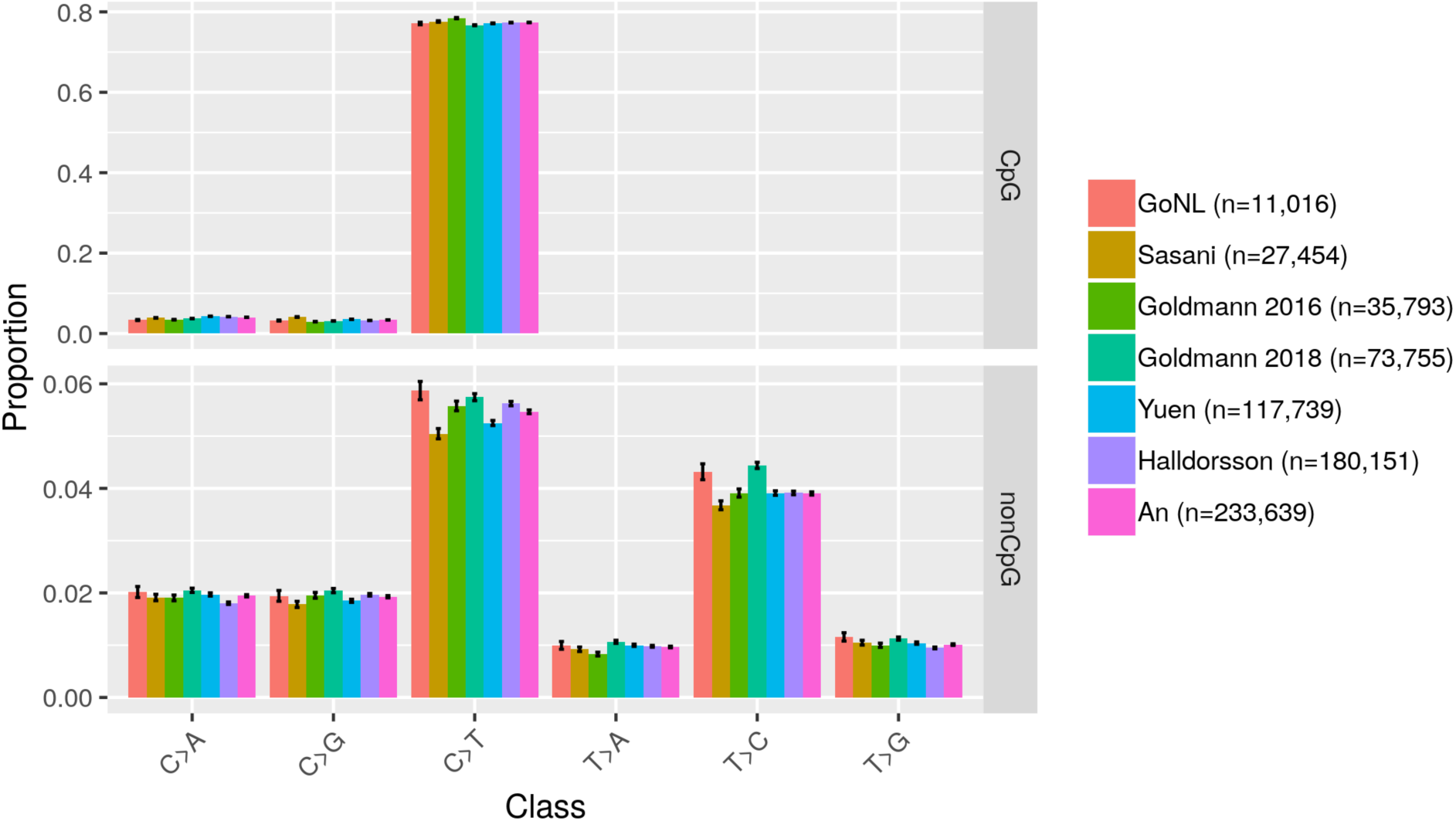
Mutation spectrum across nine mutation classes for all analyzed DNM datasets. Datasets are by order GoNL [14], Sasani et al. (2019) [41], Goldmann et al. (2016) [40], Goldmann et al. (2018) [23], Yuen et al. [39], Halldorsson et al. (2019) [38] and An et al. (2018) [42]. Error bars are binomial confidence intervals (*α* = 0.01).

**Figure S2.**
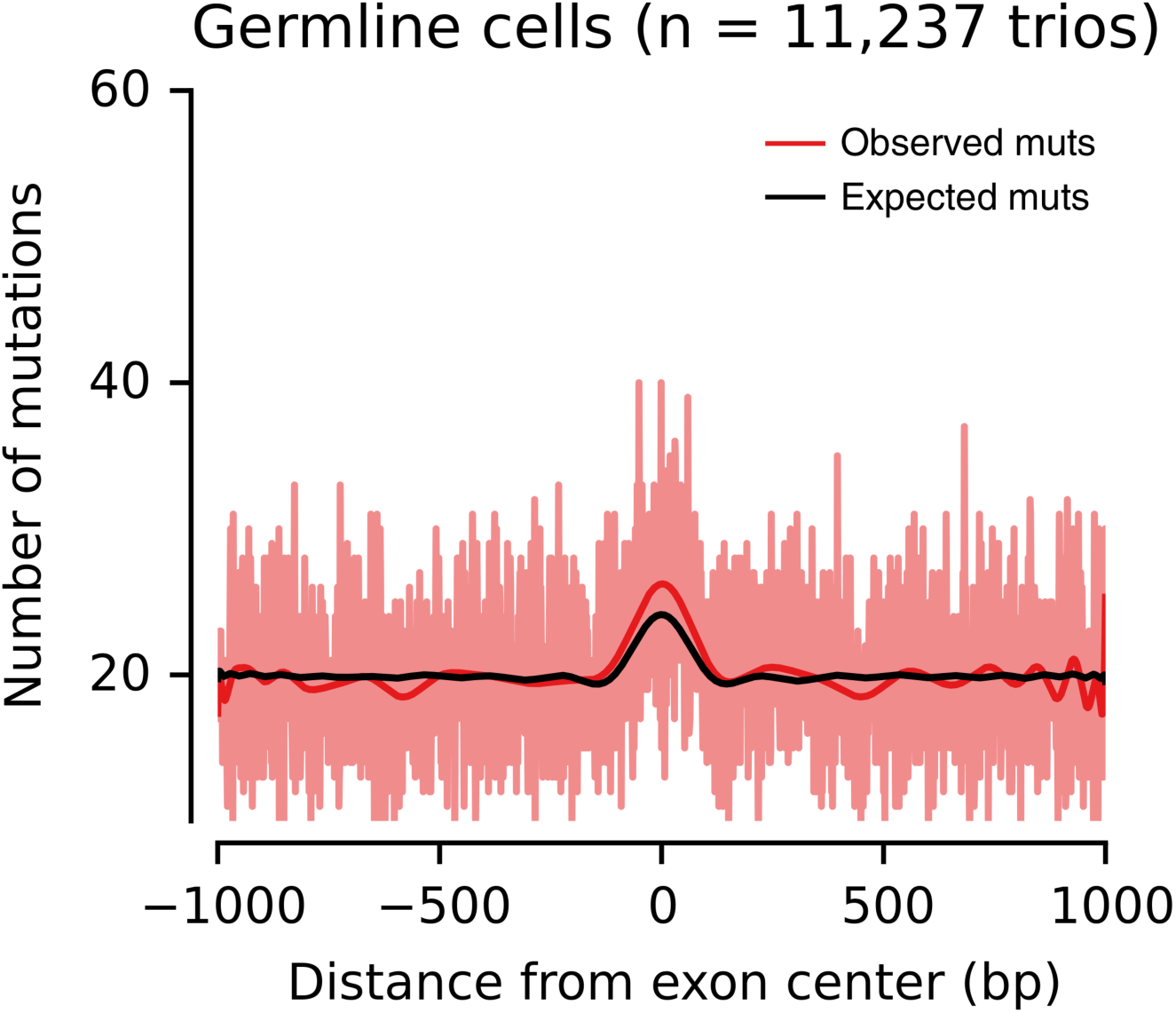
Internal exon-centered analyses on germline *de novo* mutations restricting to highly mappable regions. Exon-centered 2,001-nt-wide observed and expected mutational profiles. The light red line represents the observed number of mutations at each position, whereas the dark red and black lines represent smoothed numbers of observed and expected mutations, respectively, obtained from a polynomial fit. The observed germline exonic mutation burden is significantly increased by 6.6% (s.d. 1.7%; P=0.001, permutation-based test) compared to the expectation across introns and exons.

**Figure S3.**
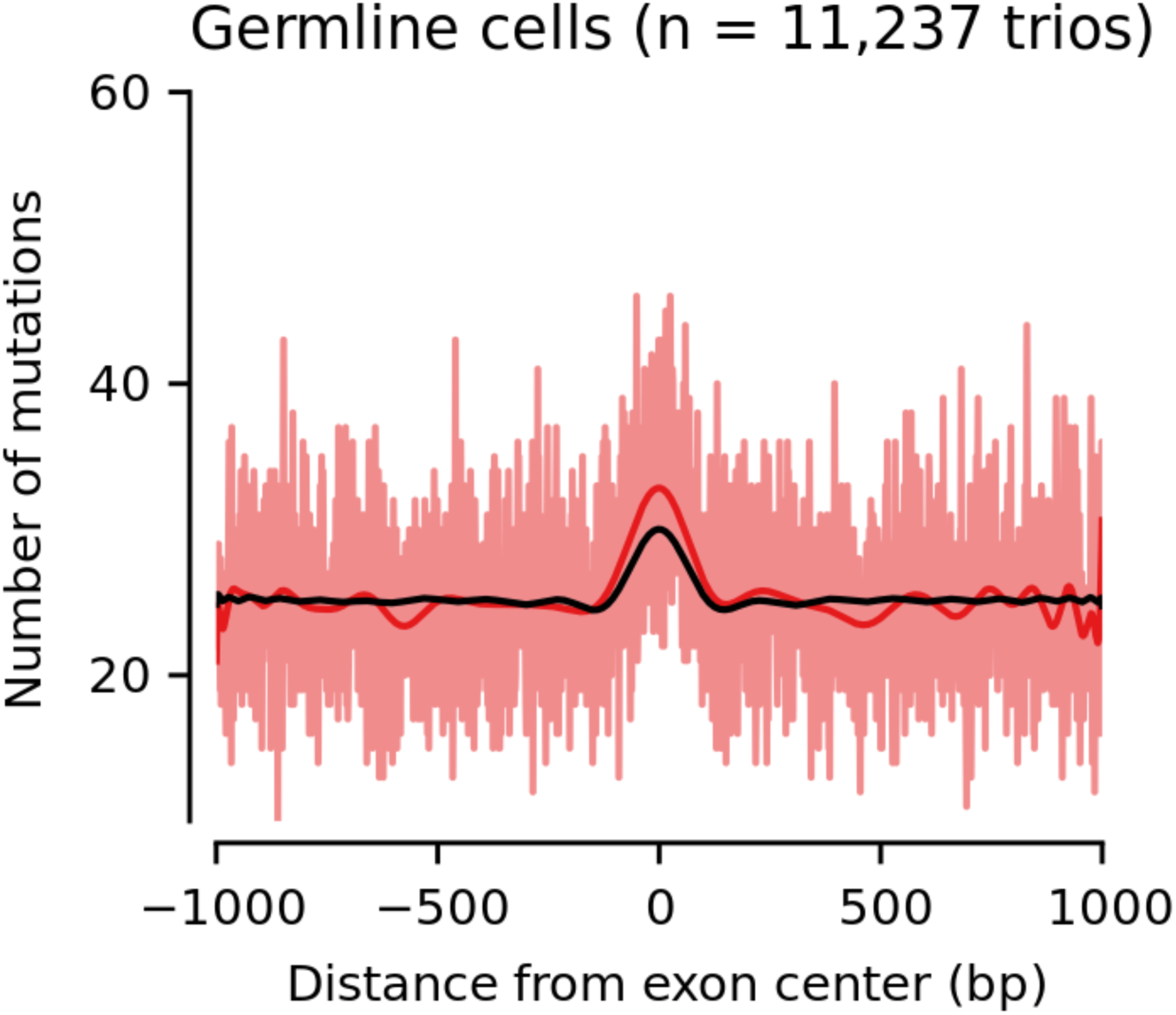
Internal exon-centered analyses on germline *de novo* mutations restricting the mutational model to genic regions. Exon-centered 2,001-nt-wide observed and expected mutational profiles. The light red line represents the observed number of mutations at each position, whereas the dark red and black lines represent smoothed numbers of observed and expected mutations, respectively, obtained from a polynomial fit. Expected was computed based on a mutational model derived only from genic regions, to test the importance of transcription-coupled repair for the mutational signature. The observed germline exonic mutation burden is significantly increased by 7.9% (s.d. 1.5%; P=0.001, permutation-based test) compared to the expectation across introns and exons.

**Figure S4.**
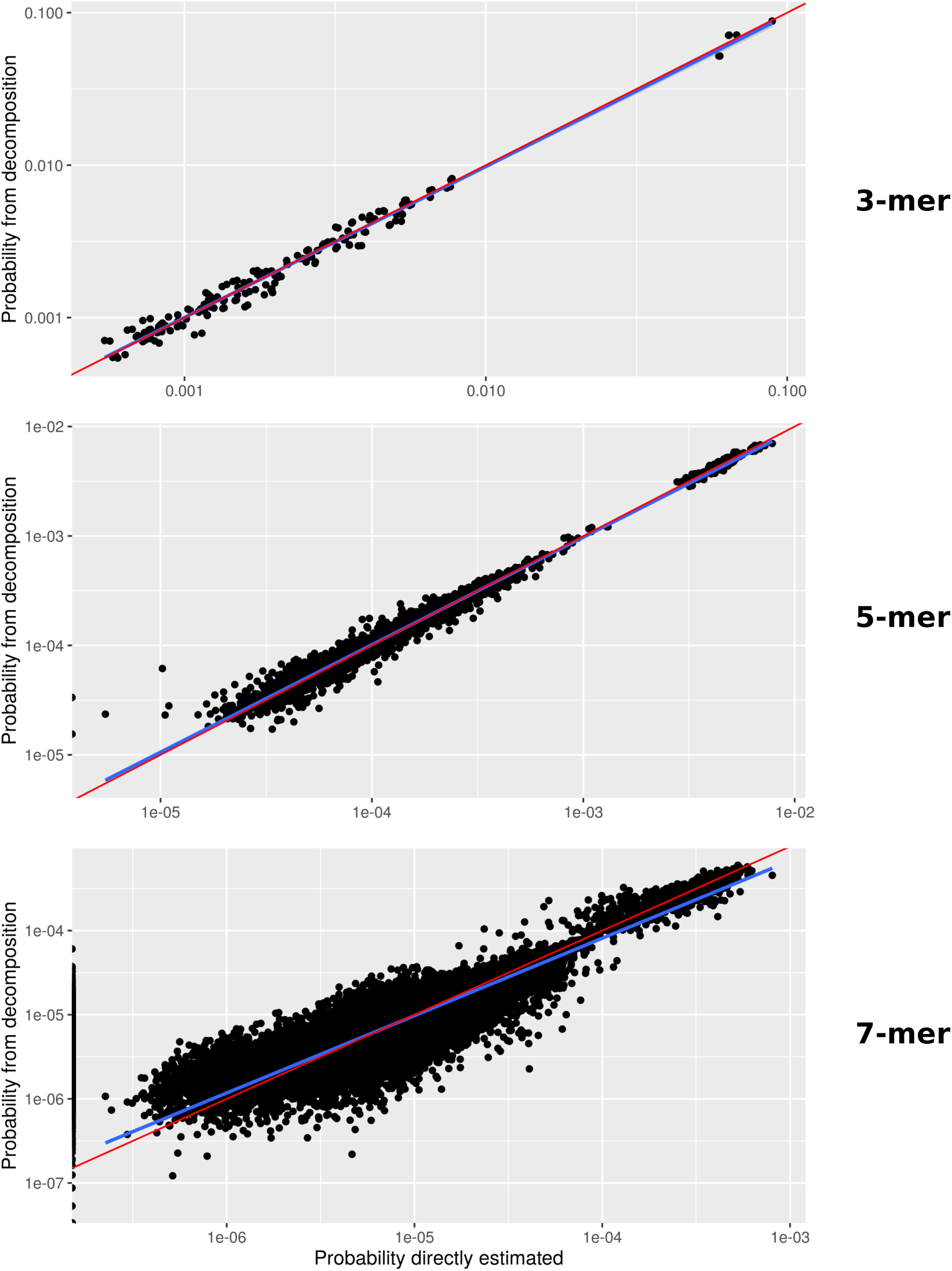
Comparison between mutation probabilities obtained from a direct vs the composite likelihood approach. Each dot corresponds to a mutation. Pearson correlation coefficients are *ρ* = 0.9966, *ρ* = 0.9979, *ρ* = 0.9858 for 3-mer, 5-mer and 7-mer, respectively. Note that the linear regression (blue line) moves further away from the diagonal (red line) as the number of values that cannot be estimated through the direct method increases (e.g. for 7-mers 1,178 out of 49,152 mutational subtypes were erroneously estimated to have a 0 probability).

**Figure S5.**
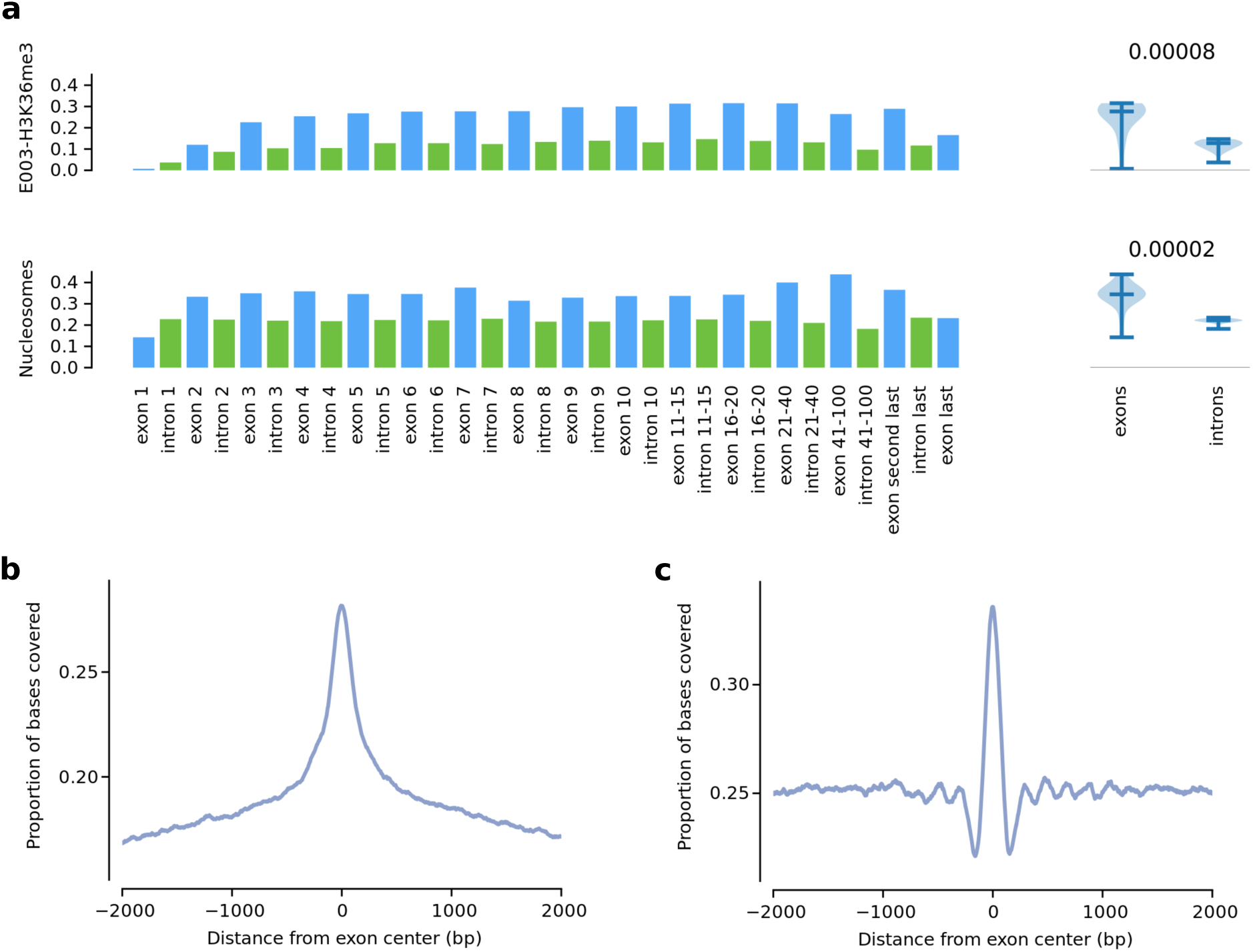
Exonic enrichment for the H3K36me3 mark and nucleosomes. Exonic and intronic coverage of H3K36me3 peaks in the H1-hESC embryonic stem cell line (E003) and of nucleosome-covered regions in GM12878 (lymphoblastoid cell line). **(a)** At the left, each bar represents the coverage of the mark in exons or introns at different positions of gene structure. At the right, violin plots show the distribution of the exonic and intronic coverage of each chromatin feature across the gene structure. The p-value from a two-tailed Mann–Whitney test compares the two distributions. Note that most of the difference comes from internal exons compared to flanking intronic sequences. **(b)** Proportion of bases covered by H3K36me3 across internal exons and flanking introns along a middle exon-centered 4001-nt window. **(c)** Proportion of bases covered by nucleosomes across internal exons and flanking introns along a middle exon-centered 4001-nt window.

**Figure S6.**
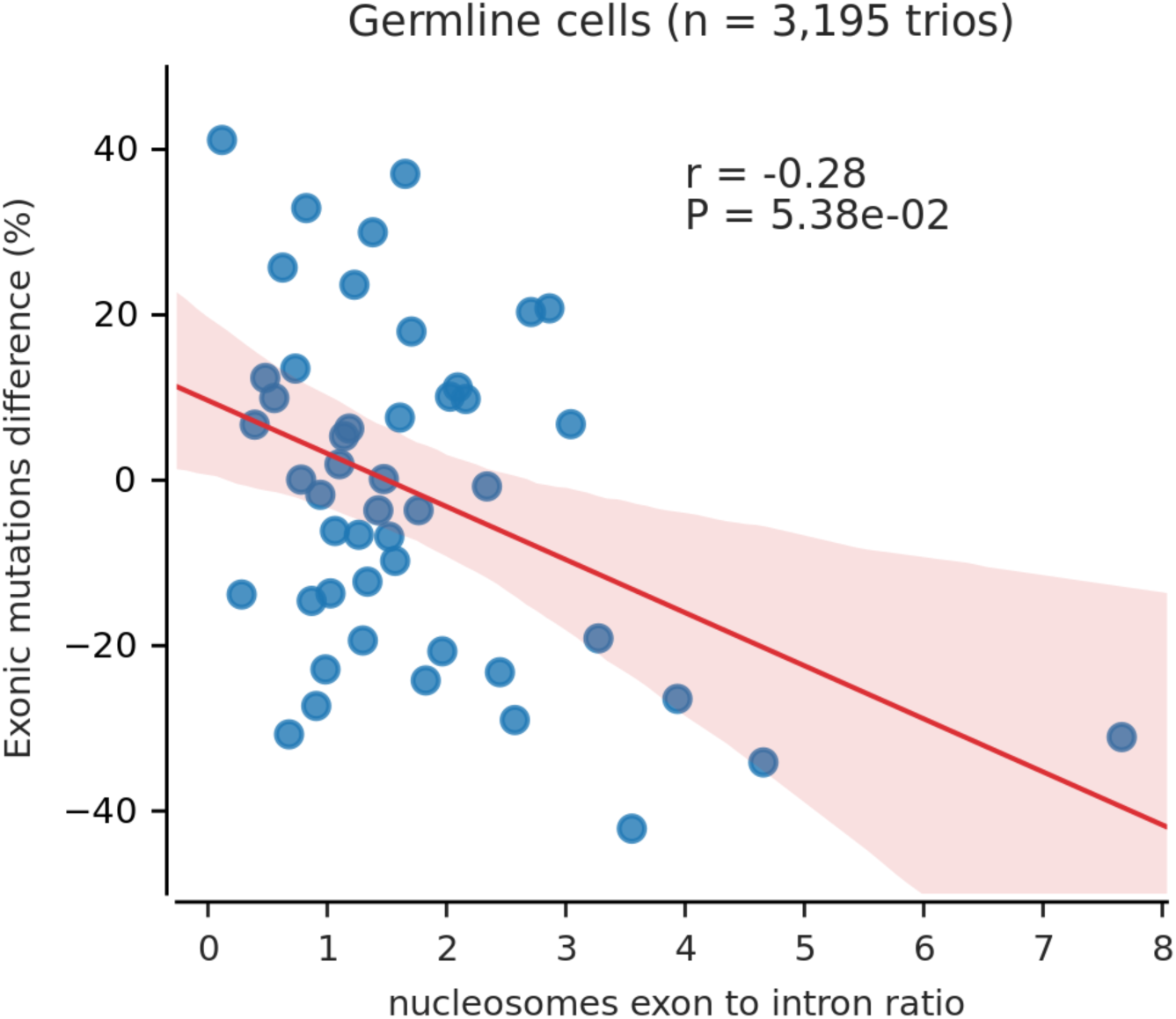
Deviation in the exonic mutation burden as a function of the nucleosome exon-to-intron ratio. Blue dots denote 50 groups of genes binned by their exon-to-intron ratio of nucleosome density (x-axis), which was derived from the lymphoblastoid cell line GM12878. The relative difference between the total observed and expected number of exonic mutations per group is shown on the y-axis. Only mutations from healthy probands were used and the expectation was computed using a 3-mer model. The trendline and its confidence interval were added using the seaborn package of Python, while the correlation coefficient and its significance were computed using an iteratively re-weighted least-squares approach as described by Frigola et al. [15].

### Supplementary Tables

**Table S1.** Proportion of sequences filtered by genome mappability issues. Middle exon-centered 2001-nt sequences with at least one affected nucleotide are classified as affected.

| Genomic regions | affected bp [%] | affected windows [%] |
| --- | --- | --- |
| UCSC blacklisted regions (low mappability) | 0.01 | 0.04 |
| hg19 low complexity (repetitive) regions | 0.68 | 22.51 |
| Intersection | 0.69 | 22.54 |

**Table S2.** Technical characteristics of the DNM datasets.

| Dataset | Sequencing technology | Mean Cov. | Calling pipeline | Quality controls |
| --- | --- | --- | --- | --- |
| GoNL [14] | Hiseq2000 | ~13x | GATK<br>(PhaseByTransmission) | a,d,e,f |
| Sasani [41] | HiseqX | ~30x | GATK | b |
| Goldmann2016 [40] | Complete Genomics | ~60x | cga-tools | a,d |
| Goldmann2018 [23] | Hiseq2000 | ~40x | GATK (joint<br>HaplotypeCaller,<br>PhaseByTransmission<br>and ReadBackPhasing) | a,c,d |
| Yuen [39]* | Hiseq2000 (n=561) | ~34x | cga-tools + GATK | d |
|  | Complete Genomics<br>(n=1,233) | ~54.5x | DenovoGear + GATK | d |
|  | HiseqX (n=3,411) | ~36.4x | DenovoGear + GATK | d |
| Halldorsson [38] | GA II, Hiseq and HiseqX | ~30x | GraphTyper | b,c |
| An [42] | Hiseq2000 (n=40),<br>HiseqX (n=1,862) | ~35.5x | TrioDeNovo +<br>DenovoGear + PlinkSeq<br>+ DenovoFlow | e |

\*Authors found no differences in DNM calling across sequencing platforms.
- (a) Random Forest classifier based on sequencing validated DNM candidates. - (b) Validation through the transmission of DNM candidates across 3 generation pedigrees. - (c) Validation through monozygotic-twin pairs. - (d) Validation through Sanger sequencing. - (e) Validation through Illumina Miseq sequencing. - (f) Validation through Ion Torrent sequencing.

**Table S3.**
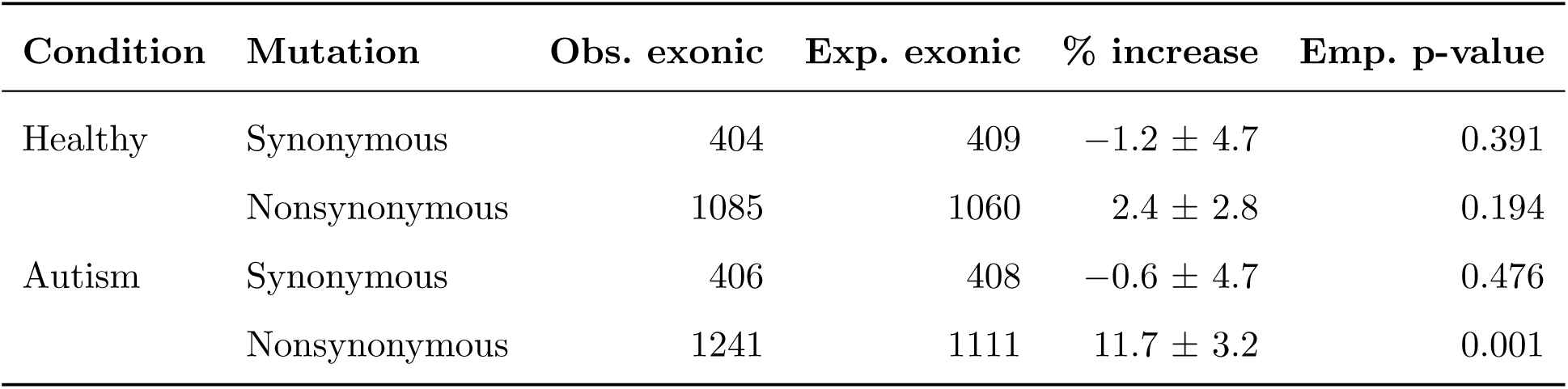
Excess in exonic burden across mutations from different conditions, stratified by mutation class. Errors denote one s.d.

| Condition | Mutation | Obs. exonic | Exp. exonic | % increase | Emp. p-value |
| --- | --- | --- | --- | --- | --- |
| Healthy | Synonymous | 404 | 409 | $-1.2 \pm 4.7$ | 0.391 |
| | Nonsynonymous | 1085 | 1060 | $2.4 \pm 2.8$ | 0.194 |
| Autism | Synonymous | 406 | 408 | $-0.6 \pm 4.7$ | 0.476 |
| | Nonsynonymous | 1241 | 1111 | $11.7 \pm 3.2$ | 0.001 |

**Table S4.**
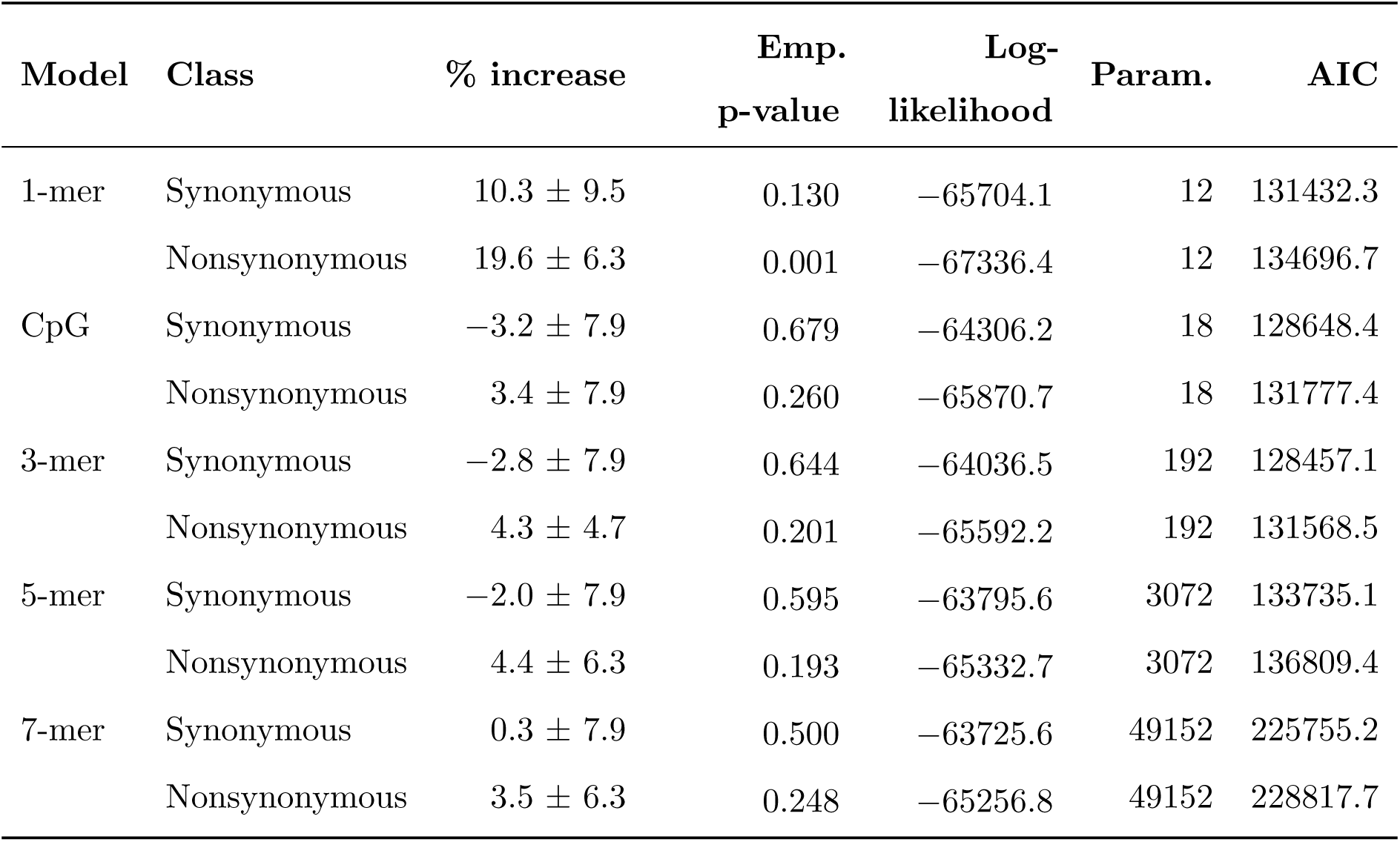
Extended sequence context dependency for Goldmann et al. (2018), stratified by mutation class. Errors denote one s.d.

| Model | Class | % increase | Emp.<br>p-value | Log-likelihood | Param. | AIC |
| --- | --- | --- | --- | --- | --- | --- |
| 1-mer | Synonymous | $10.3 \pm 9.5$ | 0.130 | -65704.1 | 12 | 131432.3 |
| | Nonsynonymous | $19.6 \pm 6.3$ | 0.001 | -67336.4 | 12 | 134696.7 |
| CpG | Synonymous | $-3.2 \pm 7.9$ | 0.679 | -64306.2 | 18 | 128648.4 |
| | Nonsynonymous | $3.4 \pm 7.9$ | 0.260 | -65870.7 | 18 | 131777.4 |
| 3-mer | Synonymous | $-2.8 \pm 7.9$ | 0.644 | -64036.5 | 192 | 128457.1 |
| | Nonsynonymous | $4.3 \pm 4.7$ | 0.201 | -65592.2 | 192 | 131568.5 |
| 5-mer | Synonymous | $-2.0 \pm 7.9$ | 0.595 | -63795.6 | 3072 | 133735.1 |
| | Nonsynonymous | $4.4 \pm 6.3$ | 0.193 | -65332.7 | 3072 | 136809.4 |
| 7-mer | Synonymous | $0.3 \pm 7.9$ | 0.500 | -63725.6 | 49152 | 225755.2 |
| | Nonsynonymous | $3.5 \pm 6.3$ | 0.248 | -65256.8 | 49152 | 228817.7 |

**Table S5.** Extended sequence context dependency for data pooled across all cohorts, stratified by mutation class. Errors denote one s.d.

| Model | Class | % increase | Emp.<br>p-value | Log-likelihood | Param. | AIC |
| --- | --- | --- | --- | --- | --- | --- |
| 1-mer | Synonymous | $13.0 \pm 3.2$ | 0.001 | -483499.6 | 12 | 967023.2 |
| | Nonsynonymous | $28.3 \pm 3.2$ | 0.001 | -495582.7 | 12 | 991189.3 |
| CpG | Synonymous | $-2.1 \pm 3.2$ | 0.786 | -468484.4 | 18 | 937004.7 |
| | Nonsynonymous | $9.8 \pm 1.6$ | 0.001 | -479643.4 | 18 | 959322.9 |
| 3-mer | Synonymous | $-1.1 \pm 3.2$ | 0.651 | -466394.2 | 192 | 933172.3 |
| | Nonsynonymous | $10.4 \pm 1.6$ | 0.001 | -477516.7 | 192 | 955417.5 |
| 5-mer | Synonymous | $-0.7 \pm 3.2$ | 0.592 | -464468.4 | 3072 | 935080.9 |
| | Nonsynonymous | $9.7 \pm 1.6$ | 0.001 | -475512.7 | 3072 | 957169.3 |
| 7-mer | Synonymous | $-0.5 \pm 3.2$ | 0.561 | -462878.9 | 49152 | 1024062.0 |
| | Nonsynonymous | $8.6 \pm 1.6$ | 0.001 | -473870.2 | 49152 | 1046044.0 |

